# Importance of viscosity contrast for the motion of erythrocytes in microcapillaries

**DOI:** 10.1101/2021.02.11.430779

**Authors:** Anil K. Dasanna, Johannes Mauer, Gerhard Gompper, Dmitry A. Fedosov

**Author notes:** Correspondence: Dmitry A. Fedosov.

## Abstract

The dynamics and deformation of red blood cells (RBCs) in microcirculation affect the flow resistance and transport properties of whole blood. One of the key properties that can alter RBC dynamics in flow is the contrast λ (or ratio) of viscosities between RBC cytosol and blood plasma. Here, we study the dependence of RBC shape and dynamics on the viscosity contrast in tube flow, using mesoscopic hydrodynamics simulations. State diagrams of different RBC dynamical states, including tumbling cells, parachutes, and tank-treading slippers, are constructed for various viscosity contrasts and wide ranges of flow rates and tube diameters (or RBC confinements). Despite similarities in the classification of RBC behavior for different viscosity contrasts, there are notable differences in the corresponding state diagrams. In particular, the region of parachutes is significantly larger for λ = 1 in comparison to λ = 5. Furthermore, the viscosity contrast strongly affects the tumbling-to-slipper transition, thus modifying the regions of occurrence of these states as a function of flow rate and RBC confinement. Also, an increase in cytosol viscosity leads to a reduction in membrane tension induced by flow stresses. Physical mechanisms that determine these differences in RBC dynamical states as a function of λ are discussed.

## 1 INTRODUCTION

Microvascular blood flow is essential for the homeostasis of organism tissues, as it transports nutrients and waste products and mediates various physiological processes. This research field has received enormous attention directed at understanding complex microvascular transport and regulation [1, 2, 3, 4, 5]. Blood is a liquid tissue whose major cellular component is erythrocytes or red blood cells (RBCs) which constitute about 45% of blood volume. A healthy RBC has a biconcave shape with a diameter of 6-8 μm and thickness of 2 μm [6]. The RBC membrane consists of a lipid bilayer and spectrin network (cytoskeleton) attached to the inside of the bilayer [7]. These structures supply cell deformability and durability, as RBCs have to frequently pass capillaries with a diameter comparable to the RBC size. The ability of RBCs to deform is vital for microvascular perfusion, as an increased membrane rigidity is generally associated with pathological conditions [8, 9] such as sickle-cell anemia [10] and malaria [11, 12].

One of the important steps toward understanding microvascular blood flow is a detailed description of RBC behavior in microcapillaries. Early experiments [13, 14, 15] have shown that RBCs passing through small vessels either deform into cup-like parachute shapes at the vessel center or assume elongated slipper shapes at an off-center position. A number of more recent microfluidic experiments [16, 17, 18, 19, 20] have systematically studied and confirmed these observations and suggested a connection between RBC elasticity and its shape in flow. From the physics point of view, it is interesting to understand how such shapes develop and which cell and flow properties determine their stability. First simple axisymmetric models of RBCs flowing in microvessels [21] have demonstrated the ability of RBCs to attain parachute and bullet-like (in very narrow vessels) shapes due to the stresses exerted by fluid flow. Two dimensional (2D) simulations of fluid vesicles mimicking RBCs have shown that cell behavior in microcapillary flow is quite complex [22, 23, 24, 25, 26, 27]. In addition to the parachute and slipper shapes, snaking dynamics (a periodic cell swinging around the tube center) at low flow rates and a region of co-existing parachutes and slippers at high flow rates were reported [24, 25]. These 2D simulations have also demonstrated that the transition between parachute and slipper shapes can be triggered by changes in flow rate or RBC membrane elasticity. This transition can be characterized by the distance between the cell’s center-of-mass and the channel center, which has been shown to have a similar behavior as a pitchfork bifurcation [23]. Nevertheless, it is still not fully clear why the parachute-to-slipper transition takes place.

Three dimensional (3D) simulations of RBCs flowing in microchannels [28, 29, 30, 31, 32, 33, 34] have confirmed the existence of stable slippers in 3D. Despite some similarities between the results obtained from 2D and 3D simulations, RBC dynamics in microchannels is inherently three dimensional, so that the results from 2D simulations are at most qualitative. For instance, 3D simulations have shown the existence of a dynamic state of RBC tumbling at a radial position away from the tube center [29, 34]. In fact, the transition from tumbling to slipper state with increasing flow rate is reminiscent of the well-known tumbling-to-tank-treading transition of RBCs in simple shear flow [35, 36, 37, 38]. Furthermore, recent experiments on RBCs in flow within square microchannels have found a tumbling trilobe state at large flow rates and low confinements [34]. Such trilobe dynamics has so far only been reproduced in simulations of RBCs in simple shear flow, and occurs at large shear rates and for large enough viscosity contrasts λ defined as the ratio between viscosities of RBC cytosol and suspending medium [39, 40], with λ ≳ 3.5.

Most of the current simulation studies assume for simplicity the viscosity contrast of unity, even though the average physiological value of λ is about five [41, 42]. The viscosity contrast is an important parameter that significantly affects RBC behavior in simple shear flow [39, 40, 43, 44, 45]. However, it remains unclear whether the viscosity contrast is important for RBC dynamics in microcapillary flow. Therefore, we focus on the effect of λ on RBC dynamical states in tube flow. Several state diagrams of RBC dynamics, including snaking, tumbling, tank-treading slipper, and parachute, are presented for different viscosity contrasts, tube diameters, and flow rates. Even though the dynamical states are similar for λ = 1 and λ = 5, there are differences in flow conditions at which they appear. In particular, the region of tumbling dynamics for λ = 5 expands toward larger flow rates in comparison to λ = 1, since an increased dissipation inside the cell suppresses membrane tank-treading in favor of tumbling motion. Furthermore, the region of parachute shapes is larger for λ =1 than that for λ = 5. A larger viscosity inside the RBC also leads to a decrease in membrane tension for the same flow conditions. Physical mechanisms that determine these differences in dynamical state diagrams for various viscosity contrasts are discussed.

## 2 MODELS & METHODS

### 2.1 Red blood cell model

A RBC is modeled as a triangulated surface with *N*_v_ = 3000 vertices, *N*_e_ edges, and *N*_f_ triangular faces [46, 28, 47, 48]. The total potential energy of the system is given by

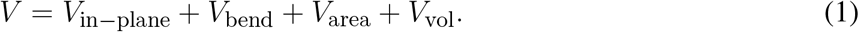

The term *V*_in–plane_ represents an in-plane elastic energy as [47, 48]

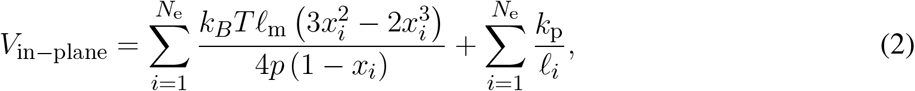

where the first term is an attractive worm-like chain potential and the second term is a repulsive potential with a strength coefficient *k*_p_. In the attractive potential, *p* is the persistence length, ℓ_*i*_ is the extension of edge *i, ℓ*_m_ is the maximum edge extension, and *x_i_* = ℓ_*i*_/ℓ_m_.

The second term in Eq. (1) corresponds to bending resistance of the membrane,

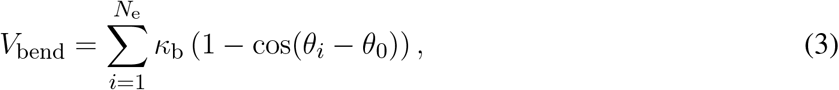

where *κ_b_* is the bending coefficient, *θ* is the angle between two neighboring faces, and *θ*_0_ is the spontaneous angle. The last two terms in Eq. (1), *V*_area_ and *V*_vol_, represent surface area and volume constraints given by

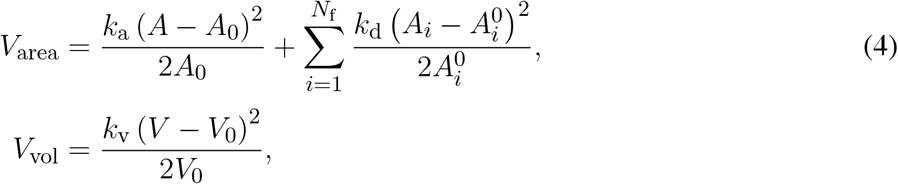

where *k*_a_, *k*_d_, and *k*_v_ are local area, total surface area and volume constraint coefficients, respectively. 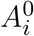, *A*_0_ and *V*_0_ are local area of individual faces, total surface area and total volume of the RBC, respectively.

### 2.2 Modeling hydrodynamic flow

The fluid is modeled by the smoothed dissipative particle dynamics (SDPD) method which is a Lagrangian discretization of Navier-Stokes equations [49, 50]. The SDPD fluid consists of *N* fluid particles which interact through conservative, dissipative and random forces. The solvent inside the RBC (cytosol) is separated from outside fluid (plasma) by the membrane. The number density of fluid particles is set to *n* =12 (per unit volume in model units) for both cytosol and plasma. Solid walls are modeled by frozen SDPD particles. Fluid-membrane interactions have two contributions: (*i*) fluid particles bounce back from the membrane surface and (*ii*) the dissipation force coefficient between fluid particles and membrane vertices is set such that no-slip boundary conditions are attained. Fluid particles are also reflected back at the solid wall. In addition, an adaptive shear force is added to fluid particles near the wall to ensure no-slip boundary conditions [51].

### 2.3 Simulation setup and parameters

Poiseuille flow with a single RBC suspended in a viscous fluid inside a cylindrical tube of length *L* = 50 *μ*m is simulated. The tube axis is aligned with the flow direction along the x axis. Diameter of the tube *D* = 2*R* determines RBC confinement as *χ* = *D_r_/D*, where 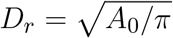 is the effective RBC diameter. To generate flow, a force *f* is applied on every solvent particle, representing a pressure gradient Δ*P/L* = *f* · *n* with the pressure drop Δ*P* along the tube length.

In simulations, cell properties correspond to average characteristics of a healthy RBC with a membrane area *A*_0_ = 133 *μ*m^2^, cell volume *V*_0_ = 93 *μ*m^3^, shear modulus *μ* = 4.8 *μ*N/m, and bending rigidity *κ* = 70 *k_B_T* =3 × 10^-19^ J [6, 52, 53, 54]. This leads to *D_r_* = 6.5 *μ*m (*D_r_* = 6.5 in model units) and a RBC reduced volume of 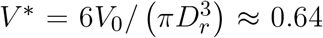. Note that the stress-free shape of a RBC elastic network [Eq. (2)] is assumed to be an oblate spheroid with a reduced volume of 0.96. Furthermore, the energy unit *k_B_T* is selected to be *k_B_T* = 0.2 in simulations, corresponding to a physiological temperature of 37° C.

To characterize different flow conditions, several non-dimensional parameters are employed

i. Reynolds number 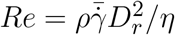 is the ratio of inertial and viscous forces, where *ρ* is the mass density, 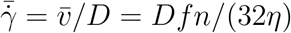 is the average (or pseudo) shear rate, and *η* is the external fluid viscosity. In all simulations, *Re* ≤ 0.3.
ii. λ = *η_i_/η_o_* is the viscosity contrast between internal (cytosol) and external (plasma) fluids. The average value of λ under physiological conditions is λ = 5 [41, 42].
iii. 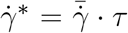 is the dimensionless shear rate that characterizes flow strength. *τ* is the RBC relaxation time given by *τ* = *ηD_r_* /*μ*.

To keep Reynolds number low enough (i.e., *Re* ≤ 0.3), in most simulations 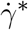 is controlled by varying η instead of changing the flow rate for a fixed viscosity.

### 2.4 Dynamical characteristics and membrane tension

To analyze dynamical properties of a flowing RBC, the gyration tensor

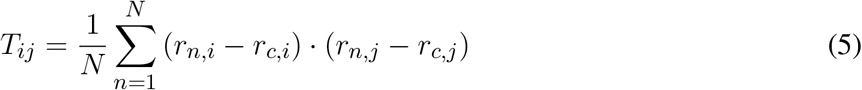

is employed, where *i* and *j* denote *x, y,* or *z*, **r**_*n*_ is the position of membrane vertex *n*, and **r**_*c*_ is the center of mass of the RBC. Then, the eigenvalues *ξ_i_* of the gyration tensor *T_ij_* characterize RBC deformation. The eigenvector that corresponds to the smallest eigenvalue is used to define the orientational axis of the cell. Orientation angle *θ*_1_ of the RBC is defined as the angle between its orientational axis and the flow direction. The eigenvalues are also used to compute cell asphericity 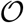, which characterizes its deviation from a spherical shape

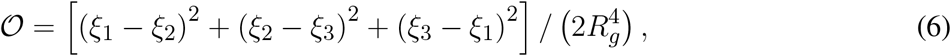

where 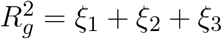.

To calculate local membrane tension *G_i_* at vertex *i*, virial stress is used as

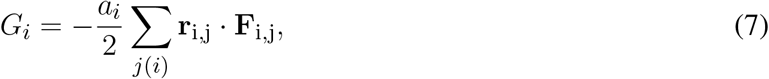

where *a_i_* is the vertex area computed as one third of a sum of all face areas adjacent to vertex *i, j*(*i*) represents all neighboring vertices connected to *i* by an edge, and **r**_ij_ and **F**_ij_ are position and force vectors at the edge (*i,j*), respectively. Note that the in-plane elastic energy, bending potential, surface area, and volume constraints can contribute to the membrane tension.

## 3 RESULTS

In microcapillary flow, RBCs are known to exhibit different dynamical states, including snaking, tumbling, tank-treading, and parachute [15, 17, 20, 24, 25, 29, 34]. Snaking is characterized by a periodic swinging in RBC orientation around the tube axis [24, 25, 29]. Tumbling is an off-axis rigid-body-like rotation, similar to RBC tumbling in simple shear flow [37, 40, 55]. Tank-treading is represented by membrane rotation with a nearly fixed cell orientation, which also occurs in simple shear flow at low enough λ [37, 40, 56]. The tank-reading state of a RBC in microcapillary flow is also often referred to as slipper. Finally, parachute is a stable stomatocyte-like RBC deformation in the tube center. These dynamical states depend on RBC mechanical properties (e.g., shear modulus, bending rigidity, viscosity contrast), cell confinement, and the flow rate. Here, we primarily focus on how the viscosity contrast affects these dynamical states for a wide range of RBC confinements and flow rates.

### 3.1 Dynamic state diagram

Figure 1 presents dynamic state diagram for the viscosity contrast λ = 5 and different *χ* and 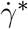 values. The representative snapshots of tumbling, tank-treading, and parachute states are also displayed (see Movies S1-S3). The snaking state exhibits minimal deformation and appears at very low shear rates 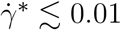 for all confinements *χ*. The tumbling state occurs for small confinements and moderate shear rates. As the shear rate increases, a tumbling RBC transits into a tank-treading state. The critical shear rate, at which the tumbling-to-tank-treading transition takes place, depends on *χ* and increases with increasing confinement. For large enough confinements and shear rates, the RBC adopts a parachute shape which exhibits least dynamics out all observed states.

**Figure 1.**
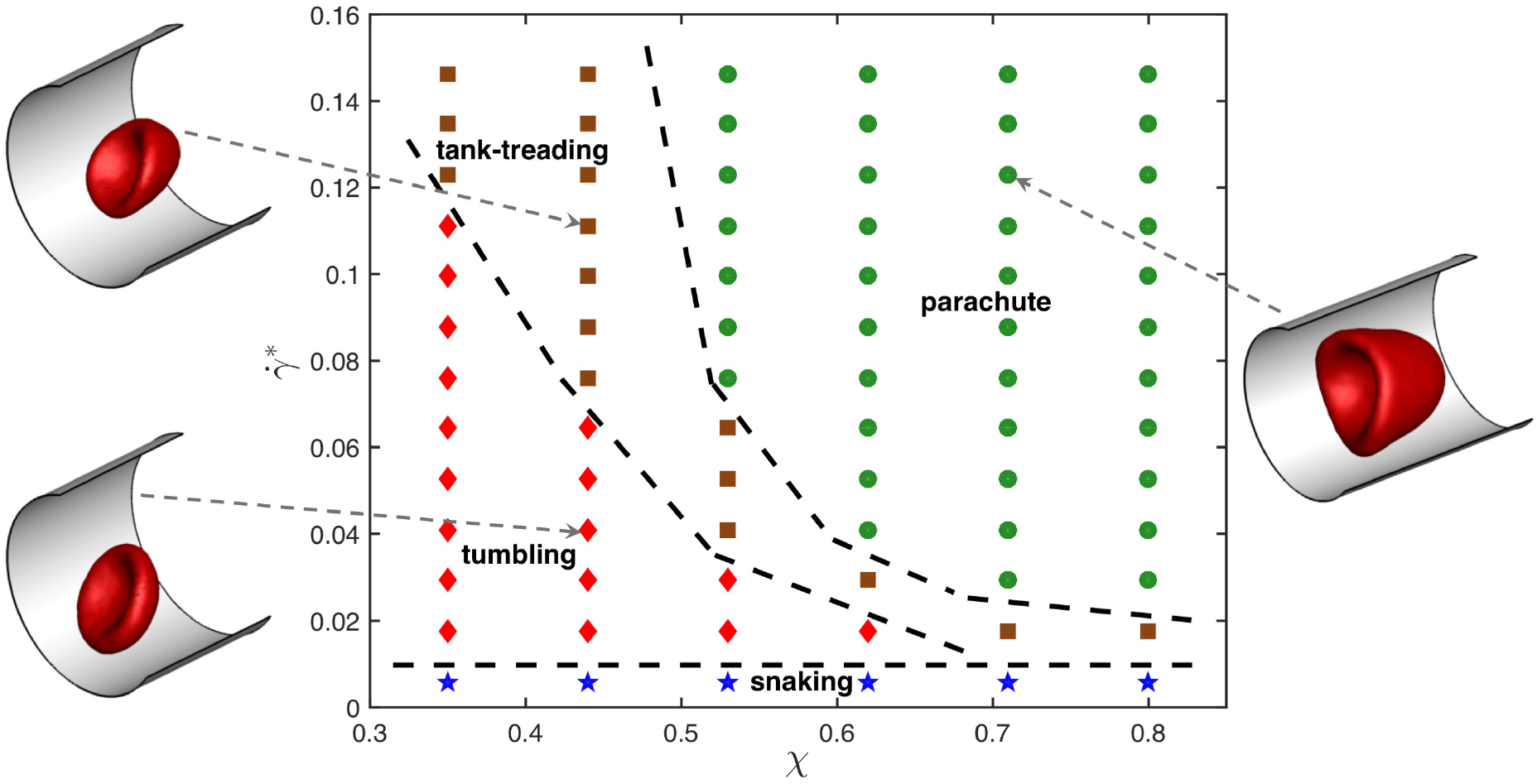
State diagram for λ = 5 showing different dynamical states of the RBC for various confinement ratios *χ* and non-dimensional shear rates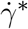. The states include snaking (blue stars), tumbling (red diamonds), tank-treading (brown squares), and parachute (green circles). Dashed lines separating regions with different states are drawn for visual guidance. Representative snapshots for tumbling, tank-treading, and parachute states are also displayed.

To understand the effect of viscosity contrast on dynamical states of the RBC in microcapillary flow, the state diagrams for λ = 1 and λ = 3 are shown for comparison in Fig. 2. As the viscosity contrast is decreased from λ = 5 to λ = 1, the parachute region widens toward smaller confinement values. This is a surprising result considering the fact that an increase in viscosity contrast suppresses tank-treading in simple shear flow [39, 40], which will be discussed later. The tumbling-to-tank-treading transition shifts toward larger shear rates as the viscosity contrast is increased from λ = 1 to λ = 5. This result is consistent with our expectations that an increase in internal viscosity leads to increased fluid stresses inside the RBC, suppressing membrane tank-treading. A similar observation has also been made in the context of adhered malaria-infected RBCs (iRBCs) under flow, such that an increase in viscosity contrast suppresses iRBC crawling at the surface and results in iRBC flipping or its complete detachment [57]. Note that the snaking state remains nearly unchanged by the viscosity contrast.

**Figure 2.**
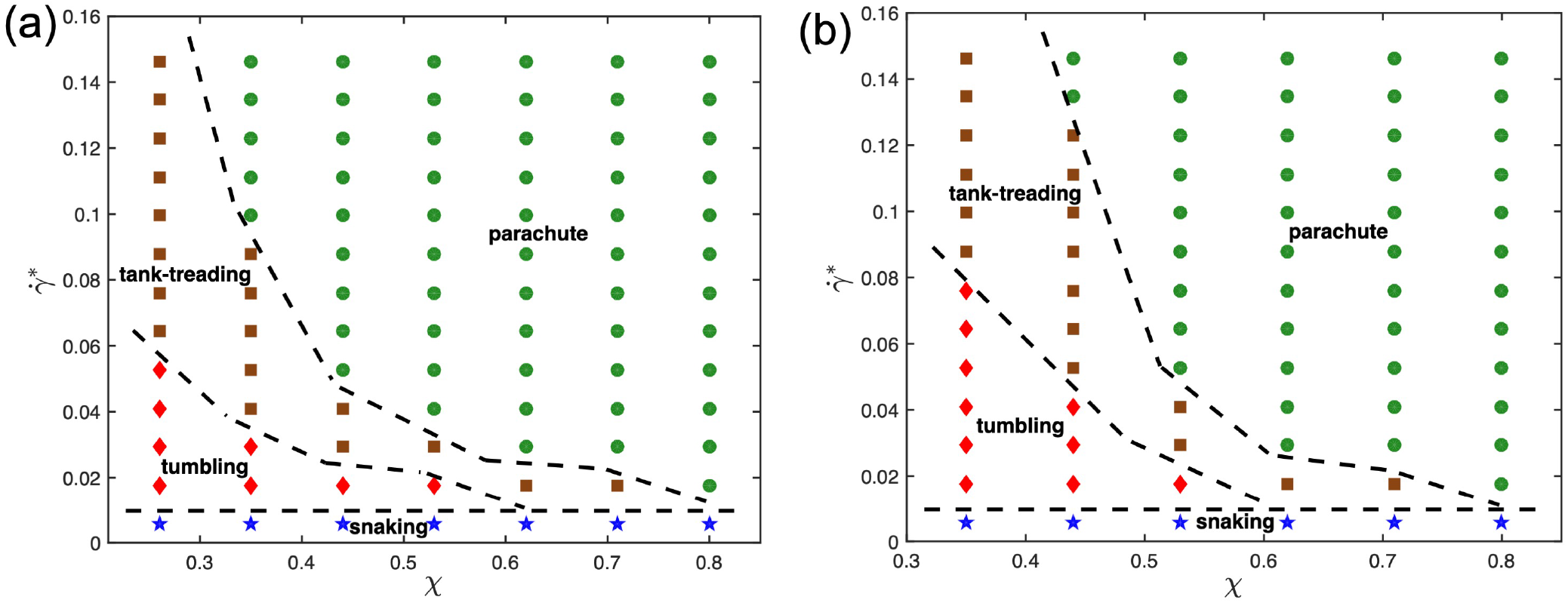
State diagrams for viscosity contrasts (a) λ =1 and (b) λ = 3 with snaking (blue stars), tumbling (red diamonds), tank-treading (brown squares), and parachute (green circles) states. Dashed lines separating regions with different states are drawn for visual guidance.

### 3.2 Dynamical characteristics

To examine differences in dynamical characteristics of RBCs with a change in viscosity contrast, multiple dynamical measures which uniquely characterize each state are computed. Figure 3 presents time evolution of the orientation angle *θ*_1_ and asphericity 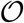 for two different flow conditions (χ = 0.35 & 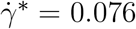; χ = 0.44 & 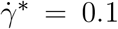) and viscosity contrasts λ = 1 and λ = 5. For the case with χ = 0.35 and 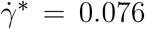 in Fig. 3(a) & (b), the RBC tank-treads for λ = 1, whereas it tumbles for λ = 5. In an idealized tank-treading state with only membrane rotation and without cell deformation, both *θ*_1_ and 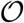 should remain constant. However, a moderate periodic deformation and oscillatory orientation swinging is observed in Fig. 3(a) for λ = 1. For the tumbling state in Fig. 3(b) with λ = 5, membrane deformation is significantly reduced, and the orientation angle spans a much wider range, indicating whole-cell flipping. For the case with χ = 0.44 and 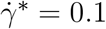 in Fig. 3(c) & (d), λ = 1 results in a parachute state with nearly constant *θ*_1_ and 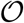, while λ = 5 leads to a tank-treading state with variations in *θ*_1_ and 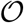 resembling those in Fig. 3(a). Interestingly, the frequency of the variations in *θ*_1_ and 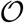 for λ = 5 in Fig. 3(d) is significantly smaller than that λ = 1 in Fig. 3(a), even though the shear rate is larger for λ = 5. This means that an increased internal viscosity slows down dynamic deformations of the RBC in microcapillary flow.

**Figure 3.**
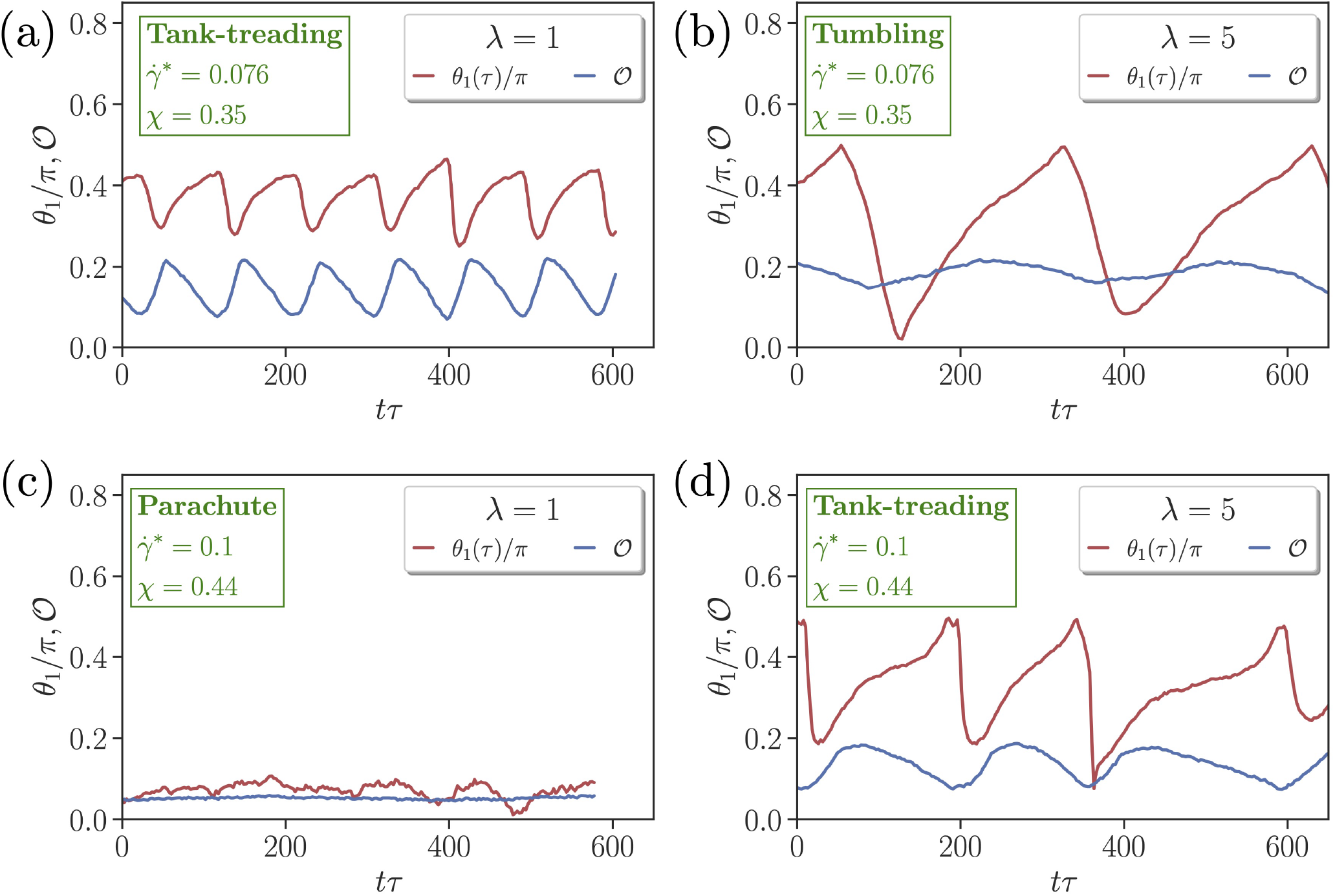
Comparison of time-dependent cell orientation *θ*_1_ and asphericity 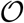 for [(a) & (c)] λ = 1 and [(b) & (d)] λ = 5. Two flow conditions are selected, including [(a) & (b)] χ = 0.35 and 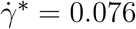, and [(c) & (d)] χ = 0.44 and 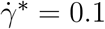. Both (a) and (d) represent tank-treading states, whereas (b) corresponds to a tumbling state and (c) to a parachute state.

### 3.3 Membrane tension

It is interesting to take a look at the effect of viscosity contrast on local membrane tension, as it might be important for the activation of mechano-sensitive channels within the membrane [58, 59]. Figure 4(a) shows the distribution of local tension *G* for a parachute shape normalized by the shear modulus *μ*. The local tension *G* is calculated using Eq. (7), where all contributions from model potentials in Eq. (1) are considered, even though the in-plane elastic-energy term supplies the maximum contribution to *G*. The concave part of the parachute shape has a significantly lower tension than the convex front of the RBC exposed to strong fluid stresses. The tension distribution for tumbling and tank-treading RBCs has a qualitatively similar trend, in which the frontal part of the cell has larger tension than the back portion. However, for tumbling and tank-treading states, local tension fluctuates in accord with the discussed RBC dynamics, while for the parachute state, temporal tension changes are generally small.

**Figure 4.**
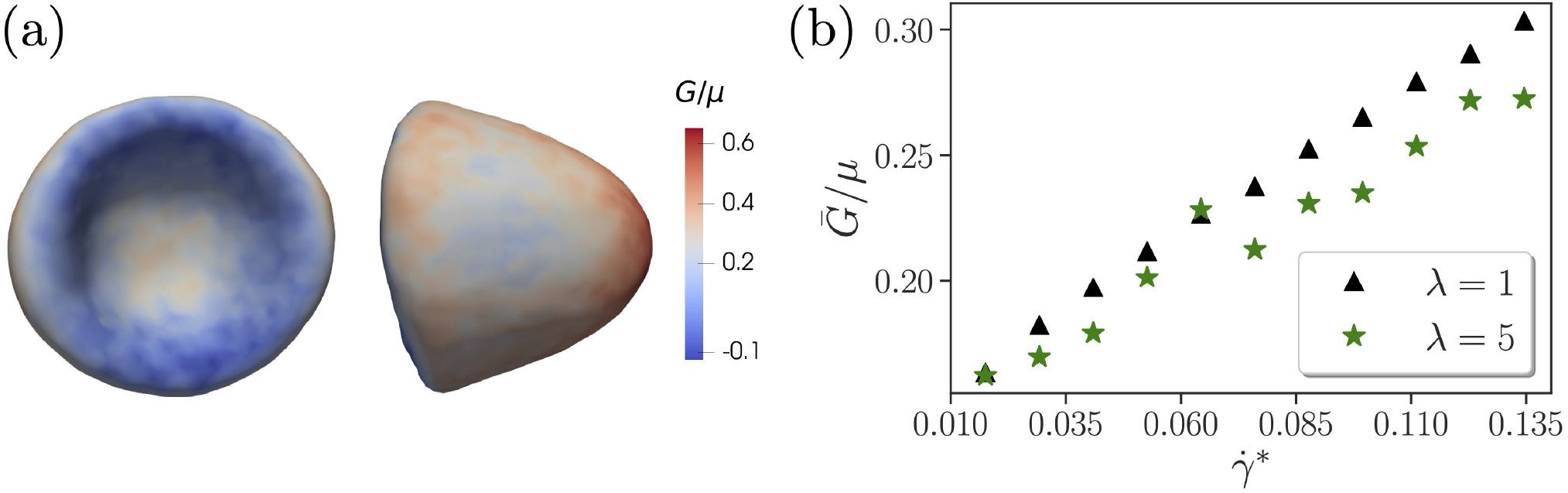
Membrane tension. (a) Side and front views of the parachute shape with a local tension *G* indicated by the color code and normalized by the shear modulus *μ*. Here, λ = 5, χ = 0.71, and 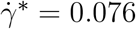. (b) Average tension 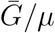 of the whole RBC as a function of non-dimensional shear rate 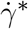. Here, the confinement is fixed at χ = 0.53. The data are shown for two different viscosity contrasts λ =1 and λ = 5. A jump in tension for λ = 5 at 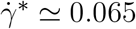 corresponds to the tank-treading-to-parachute transition.

Figure 4(b) presents tension 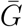 averaged over the RBC surface as a function of 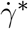 for two different viscosity contrasts λ =1 and λ = 5. The average tension increases with the shear rate 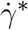 in an almost linear fashion for both viscosity contrasts. Interestingly, λ = 5 generally leads to a lower tension in comparison with λ =1, which is consistent with a previous numerical investigation [60] showing that the maximum tension increases with the flow rate and decreases with increasing viscosity contrast. For λ = 5, there is a jump in tension at approximately 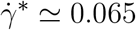 that corresponds to the tank-treading-to-parachute transition as shown in Fig. 1. Note that such jump is not present for λ = 1.

An average tension of about 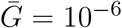 N/m (or 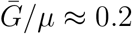) is comparatively large. For example, in a recent study on sculpting of lipid vesicles by enclosed active particles [61], complex vesicle shapes have been observed for floppy vesicles with a tension of about 10^-8^ N/m, while a high membrane tension of about 10^-5^ N/m completely suppresses any vesicle shape changes. Apart from the average tension, it is also instructive to look at the maximum tension *G*_max_ = max{*G_i_*} for different χ and 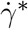. As expected, *G*_max_ increases with increasing shear rate. For λ = 5, the maximum tension at χ = 0.62 is *G*_max_/*μ* = 0.7 for 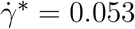 and *G*_max_/*μ* = 0.77 for 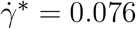 (both are parachute states). For a given shear rate, an increase in confinement results in elevation of *G*_max_, e.g. for 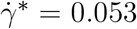, *G*_max_/*μ* = 0.64 for χ = 0.35 (tumbling state) and *G*_max_/*μ* = 0.7 for χ = 0.62 (parachute state). These trends are similar for λ =1. However, differences in Gmax with respect to the viscosity contrast are rather small, indicating that external fluid stresses mainly govern the membrane tension. The magnitudes of maximal tension from our simulations are consistent with the values reported in Ref. [60].

## 4 DISCUSSION AND CONCLUSIONS

In our study, we have focused on the effect of viscosity contrast λ on RBC dynamic states in microcapillary flow. State diagrams with different dynamic states, such as snaking, tumbling, tank-treading, and parachute, have been constructed for several λ values and wide ranges of non-dimensional shear rates 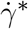 and confinements χ. Our central result is that there are significant changes in the state diagram when the viscosity contrast is decreased from λ = 5 to λ = 1. In particular, the region of stable parachutes becomes larger and expands toward lower confinements with decreasing λ. This result seems to be in contradiction to the fact that a large viscosity inside the RBC dampens membrane dynamics and hence, should suppress the dynamic tank-treading state [39, 40]. To verify the robustness of our simulation predictions, we have performed simulations with a consecutive change in the viscosity contrast for several conditions where parachutes are stable for λ = 1 and tank-treading is stable for λ = 5. Thus, after reaching a stable parachute state for λ =1, the viscosity contrast is instantaneously switched to λ = 5, leading to the tank-treading state. Then, switching back to λ =1 brings the initially tank-treading RBC to the parachute state. Furthermore, a larger parachute region for λ =1 than that for λ = 5 has also been observed in 2D simulations of vesicles [24, 25].

To reconcile this seeming contradiction, physical mechanisms that govern the parachute-to-tank-treading transition in tube flow have to be uncovered. A study based on 2D simulations of vesicles [23] suggests that the parachute-to-slipper transition can be described well by a pitchfork bifurcation and that a slipper shape provides a higher flow efficiency for RBCs. Unfortunately, this argument has not been connected in any way to the viscosity contrast or internal cell dissipation. From existing experimental and simulation studies [15, 17, 20, 24, 25, 29, 34], it is clear that the parachute state requires large enough flow rates, such that flow stresses in the tube center are sufficient to deform the RBC into a parachute shape. Therefore, only when a RBC conforms well enough to the flow profile, the parachute state is stable. Note that we have not found substantial differences in parachute shapes for different viscosity contrasts, which is likely due to the fact that a non-dynamic parachute state of the RBC depends primarily on its elastic properties, and is nearly independent of internal dissipation. Our hypothesis is that membrane dynamics is important for parachute stability, in the following sense. As the parachute-to-tank-treading transition is approached, a perturbation (e.g., due to cell diffusion) in RBC position from the tube center leads to asymmetry in fluid-flow stresses which pull the RBC away from the center and set the membrane into a tank-treading-like motion. A slight motion of the membrane in the parachute state is observed in our simulations, as the RBC is never perfectly symmetric and is often located slightly away from the tube center. For λ = 1, the membrane can rotate faster than in case of λ = 5, and therefore, the mismatch between local membrane motion and fluid flow is smaller, resulting in reduced local fluid stresses that pull the RBC away from the center. For λ = 5, the local fluid stresses on the RBC are larger due to slow membrane tank-treading, leading to the destabilization of parachute shape at larger confinements in comparison to λ = 1.

Another important difference in the state diagrams for λ = 1 and λ = 5 is that the tumbling-to-tank-treading transition occurs at larger shear rates for λ = 5 than for λ = 1. This can be explained by the fact that an increased dissipation inside the RBC for λ = 5 suppresses tank-treading motion and delays the transition in terms of 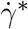. In fact, in simple shear flow, the tank-treading state does not exist for λ = 5 [39, 40]. For microcapillary flow, RBC tank-treading becomes possible at λ = 5 due to the confinement which can trigger the tumbling-to-tank-treading transition even when cell dimensions are smaller than the distance between two walls [62]. For a large enough vessel diameter, it is plausible to expect that the tank-treading state should disappear for λ = 5, as local flow conditions should closely resemble simple shear flow at the scale of RBC size. For instance, recent microfluidic experiments in a square channel [34] have reported the existence of rotating trilobe shapes at low confinements and high flow rates, which are consistent with RBC shapes in simple shear flow at λ = 5 [39, 40].

Membrane tension must be directly related to mechano-transduction as RBC membrane contains many mechano-sensitive channels [58, 59]. We have shown that an increase in the viscosity contrast lowers the membrane tension. A high viscosity of the cytosol provides a large dissipation, reducing membrane tension. Furthermore, the maximum tension increases with increasing shear rate 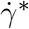 and confinement χ. Several experimental studies show that flow stresses can change RBC biochemical properties. For instance, when RBCs pass through small constrictions, they release ATP which can participate in vasodilation signaling [63, 64]. Furthermore, a recent investigation [65] reports that when RBCs pass through small constrictions, the mechano-sensitive channels (e.g., Piezo1 and Gardos channels) that participate in RBC volume control become activated. The relevance of membrane tension has also been demonstrated for malaria disease, such that an increased RBC membrane tension in Dantu blood group significantly reduces the invasion of RBCs by malaria parasites, which is a protective mechanism from malaria infection [66].

## CONFLICT OF INTEREST STATEMENT

The authors declare no conflicts of interest.

## AUTHOR CONTRIBUTIONS

A.K.D and J.M. performed simulations and analyzed the data. G.G. and D.A.F. designed research. D.A.F. supervised the project. All authors discussed the results and wrote the manuscript.

## Notes

### Competing Interest Statement

The authors have declared no competing interest.

